# Overlap of movement planning and movement execution reduces reaction time by up to 100ms

**DOI:** 10.1101/039842

**Authors:** Jean-Jacques Orban de Xivry, Valéry Legrain, Philippe Lefèvre

## Abstract

Motor planning is the process of preparing the appropriate motor commands in order to achieve a goal. This process has been largely considered as occurring before movement onset and has been traditionally associated with reaction time. However, in a virtual line bisection task, we observed an overlap between movement planning and execution.

In this task performed with a robotic manipulandum, we observed that the participants (N=30) made straight movements when the line was in front of them (near target) but made often curved movements towards a farther target that was located sideways in such a way that they crossed the line perpendicular to it. Unexpectedly, movements to the far targets had shorter reaction times than movements to the near target (mean difference: 32ms, SE: 5ms, max: 104ms). In addition, the curvature of the movement modulated reaction time. A larger increase in movement curvature from the near to the far target was associated with a larger reduction in reaction time. These highly curved movements started with a transport phase during which accuracy demands were not taken into account.

We concluded that accuracy demand imposes a reaction time penalty if it is processed before movement onset. This penalty is reduced if the start of the movement can consist of a transport phase and if the movement plan can be refined in function of accuracy demands later in the movement, hence demonstrating an overlap between movement planning and execution.

**New and Noteworthy:** In the planning of a movement, the brain has the opportunity to delay the incorporation of accuracy requirements on the motor plan in order to reduce the reaction time by up to 100ms. Such shortening of reaction time is observed here when the first phase of the movement consists in a transport phase. This forces us to reconsider the idea that motor plans are fully characterized before movement onset.

## Introduction

Motor planning is the process of selecting a goal and the appropriate motor commands in order to achieve this goal. This process has been largely considered as part of a single building block that specifies all the characteristics of the ensuing movement. For instance, the latest theories of motor planning suggest that the "complete specification of the motor command" occurs before movement start (Wong et al., 2015). Optimal control theory (Harris and Wolpert, 1998; Todorov and Jordan, 2002; Scott, 2004) also suggests that the control policy is defined before movement onset. In case of multiple possible motor plans, the average of these control policies is used early on and the appropriate motor plan is selected later (Stewart et al., 2014; Gallivan et al., 2016a, 2016b).

The idea that movement should be completely pre-planned before its execution is reminiscent of the work of Henry and Rogers in the 60’s (Henry and Rogers, 1960). In their memory drum theory of movement preparation, they suggested that planning took longer for more complex movements, which was reflected in the movement reaction time (RT). More precisely, Henry and Rogers (1960) found that reaction time was shorter for simply lifting a finger than for reaching to a goal with a single movement or with a more complicated sequence of movements. However, in these experiments, distance to the target or accuracy demands (i.e. how accurate you have to be, which depends on target size etc.) are confounded with movement complexity. These factors nonetheless influence movement reaction time. For instance, Laszlo and Livesey (1977) found that reaction times were 100ms shorter for non-goal directed movements (when accuracy requirements were removed) than for goal directed movements with strict accuracy demands (while movement complexity was matched). In contrast, loosening accuracy requirements by making the width of the target larger does not influence reaction time (Quinn et al., 1980; Orban de Xivry, 2013). Considering that non-goal directed movements are movements with very loose accuracy requirements, there seems to be a conflict between the effect of loosening or completely removing accuracy demands on reaction time (only the latter influences reaction time). The aim of this paper is to resolve this apparent contradiction by looking at the actual influence of accuracy demands on movement reaction time.

To do so, we set up a line bisection task in which the position of the lines forced the participants to adopt different strategies for different conditions but accuracy demands were identical across conditions. As a result, we observe that when the movement began with a transport phase (requiring less accuracy), the reaction time was much shorter than when the participants moved directly towards the center of the line. In other words, delaying the influence of accuracy demands on movement kinematics (because of the transport phase) led to shorter reaction times, which were up to 100ms shorter than in the control condition. That is, delaying the planning for movement accuracy during execution led to a substantial decrease in reaction time. These data suggest that increasing the overlap between movement planning and movement execution leads to a reduction in reaction time.

## Methods

### Participants

Thirty one healthy young participants were recruited to participate in our experiment. All participants had no known history of neurological disorders and no recent trauma of the upper-limbs or residual consequence of it, were right-handed and between 20 and 38 years old (mean: 24 years), and normal or corrected-to-normal vision. All of them gave written informed consent. The procedures were approved by the Université catholique de Louvain Ethics Committee and were in accordance with the latest version of the Declaration of Helsinki.

### Experimental setup

Participants sat in front of a robotic arm (Endpoint Kinarm, BKin Technologies, Kingston, Ontario, Canada). They controlled the handle of the robot in order to move a white cursor (disk with diameter of 0.5cm) that was displayed on a horizontal mirror positioned above the arm. The cursor and targets of interest were displayed on a screen placed tangentially above the mirror and were reflected by it. Because the mirror was halfway between the handle and the screen, the cursor appeared to be positioned at the same position in space as the hand after horizontal positions were properly calibrated. With this setup, subjects could not see their hand and the displayed cursor was the only available visual feedback of their arm position.

The robot controlled the display through custom-made MATLAB (R2007) programs uploaded to a real-time computer. It also monitored hand position, velocity and acceleration at 1000Hz. Kinematic and dynamic data were stored on a PC for offline analysis.

### Protocol

In this experiment, participants were instructed to perform a line bisection task by moving the cursor towards lines projected on the horizontal mirror and crossing them at their middle point (Fig.1). At the start of each trial, participants were required to bring the hand cursor inside a 2cm x 2cm orange square that was located on the lower right part of the workspace. When the cursor was inside the square, an orange line appeared at one of four possible positions. There were three possible line lengths (10, 15 or 20cm). The participants were instructed to drive the cursor through the middle of the line with a continuous and smooth movement. For each position, the line was presented either horizontally or vertically (see Fig.1). The center of horizontal lines was either 5cm or 22.5cm on the left of the starting point and 10cm above it. The center of the vertical lines was 10cm on the left of the starting point and either 5cm or 22.5cm above it. With this design, each line position was associated with only one line orientation. Lines were presented after a random time interval once the hand cursor was inside the starting position (1000-1500ms) so that the participants could never anticipate where and when the next line would appear. Once the participants had crossed the line, they were required to go back to the starting position and the next trial was initiated after an inter-trial time interval of 200ms. Participants were instructed to move at a comfortable speed. Each block consisted of 5 sub-blocks of 12 trials (2 line orientations x 2 distances x 3 lengths) that were randomly presented. Participants performed two blocks for a total of 120 trials.

**Figure 1:**
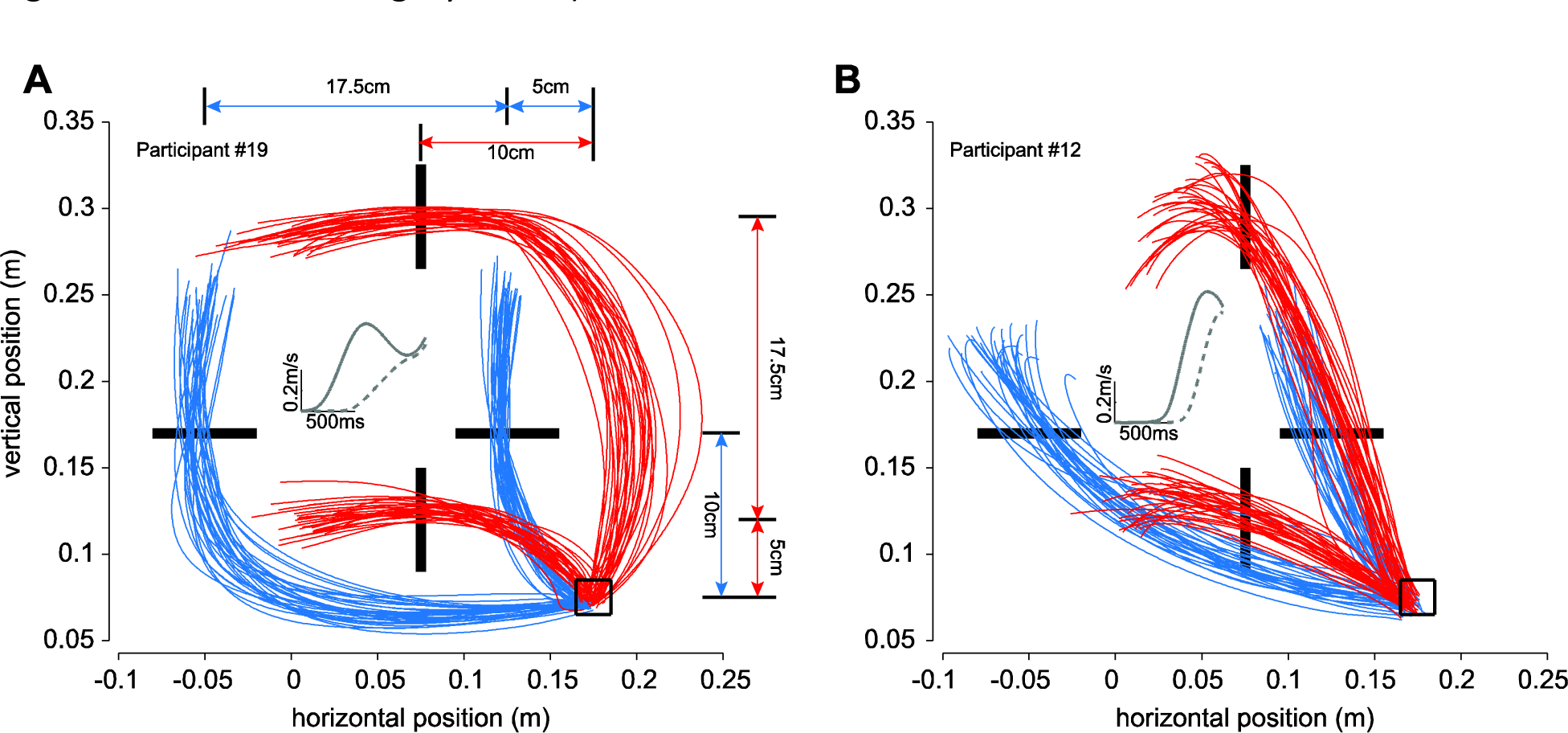
Illustration of hand trajectories from two typical subjects. For each subject, the starting position is represented by the square and the target lines are represented as black bars. For clarity, the lines are shorter in the illustration than in reality (10, 15 or 20cm). Hand trajectories of all the trials for these participants are given for horizontal lines (blue traces) and for vertical lines (red traces). The distance between the different targets and the starting position are given in panel A. For each subject, the inset in the middle of the figure represents the average vectorial velocity trace (aligned on time of reaching to the target) for horizontal near and far targets (resp. dotted and solid lines).

Movement curvature was measured as the average distance between the actual hand trajectory and a straight line linking movement onset and offset. Another dependent measure was reaction time (RT, time between the line appearance and the start of the movement), which corresponded to the time at which the vectorial velocity reached 2cm/s. Trials in which movement onset was detected before target appearance or more than 750ms after target appearance were rejected from the analysis (8% of the trials). Movement duration corresponds to the time between movement onset and when the hand cursor crossed the line. The absolute position error was computed as the distance between the hand cursor and the middle of the line at the time of line bisection. Finally, to quantify movement variability of these 2D movements, we first normalized the time of movement (0% is the movement onset and 100% is line bisection) and resampled them in 0.4% time intervals (250 samples) using spline functions (as in Orban de Xivry et al., 2011). For each of the 250 samples, we computed the minimum distance between the 2D cursor position and the average movement trajectory. Across-movement variability was equal to the average distance across movements. This measure of variability was normalized by its value on the first sample (at movement onset) in order to account for inter-subject variability.

Line length was used only to avoid that the participants would always reach to the exact same point but its influence on the behavior was not analyzed. Data were therefore merged together according to the remaining conditions.

For each participant, median reaction time (instead of mean given the skewness of reaction time data) and mean curvature, absolute position error, movement duration and normalized variability were computed. They were used as dependent variable in two repeated-measure ANOVAs with 2 within-subject factors: *orientation of the line* (horizontal or vertical) and *distance of the line* (near or far). Tukey’s post-hoc tests were used for one-to-one comparisons. Changes in the variables of interest were obtained by subtracting their value for the near target from the same measure for the far target independently for each line orientation.

Statistical tests were performed with Statistica (Dell Inc.). Level of significance was 0.05. Effect sizes were evaluated through partial eta-square. The effect of period was never found significant, neither as main effect nor in an interaction. This effect is thus not reported in the Results section.

One participant had to be excluded from the analysis because he failed to comply with the task instructions and did not aim at the center of the lines.

## Results

In this experiment, we asked participants to bisect lines of different length with a cursor. In this task, the directionality of the target led to a wide between-participant variety of hand paths. This variety of strategies is illustrated in Fig.1 for two participants. The first participant presented in Fig.1 (participant #19, panel A) exhibited simple and slightly curved movements towards the near targets. The velocity profile of these movements to the near targets exhibited a single peak. In contrast, the movements to the far targets were highly curved and the corresponding average velocity profiles exhibited a local minimum (inset of Fig.1A, solid grey trace). In other words, this participant chose a curved hand trajectory in such a way that it crossed the line perpendicular to it. However, there was a lot of variability across participants in terms of movement curvature. Some participants, as the one illustrated at Fig.1B, did not exhibit highly curved movements for the far targets and the velocity profiles for both targets only exhibited a single peak (inset of Fig.1B, solid and dotted grey traces).

This feature of the movements was characterized by the movement curvature, which was much higher for movements to the far than to the near targets (main effect of target distance: F(1,29)=150.69, p<0.0001, partial eta-square: 0.84, Figure 2A). In addition, the change in movement curvature seems slightly larger for vertical than horizontal lines (interaction between orientation and distance: F(1,29)=5.94, p=.02, partial eta-square: 0.17). Nonetheless, the effect of target distance on movement curvature was highly significant for both line orientations (Tukey’s post-hoc tests: p<0.0002 for both).

**Figure 2:**
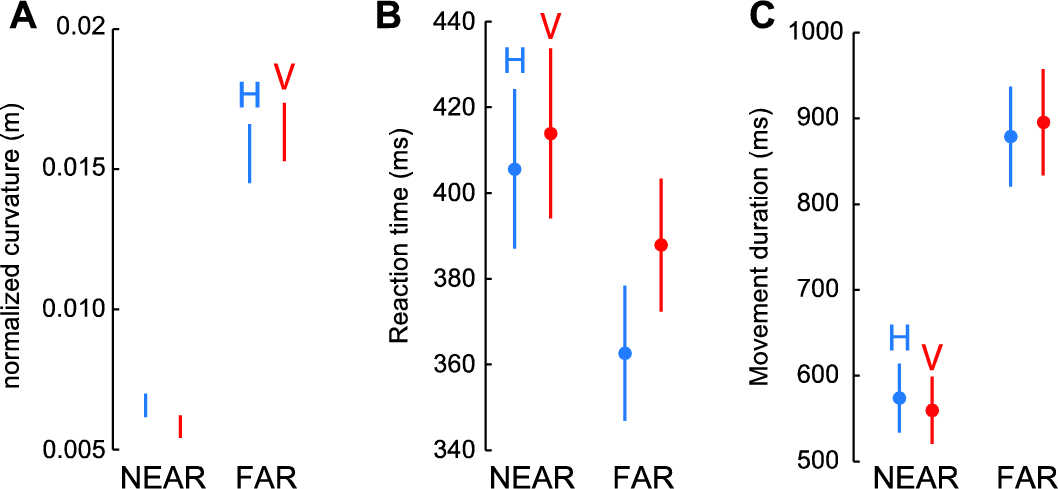
Influence of line orientation and distance on normalized movement curvature (panel A), on reaction time (panel B) and on movement duration (panel C). Blue is associated with horizontal lines (H) and red with vertical lines (V). Error bars represent standard error of the mean.

Despite being more curved, movements to the far target had shorter reaction times than movements to the near target (Figure 2B). This effect gave rise to a main effect of target distance on reaction time (F(1,29)=46.36, p<.00001, partial eta-square: 0.61) but also to an interaction between line direction and target distance (F(1,29)=8.62, p=.006, partial eta-square: 0.23), which was due to the fact that the decrease in reaction time with target distance was more pronounced for horizontal lines than for vertical lines. However, this effect remained significant for both line orientations (Post-hoc test: p<0.0002 for both orientations; ΔRT H: 40ms±5.8ms (mean±SE); ΔRT V: 24ms±5.5ms). That is, it took less time for the brain to prepare a more curved movement towards a farther target. Nonetheless, accuracy to near and far targets was essentially identical (absolute position error: main effect of target distance: F(1,29)=0.55, p=0.46, partial eta-square: 0.018). However, this change in reaction time was insufficient to compensate for the longer movement duration (from movement onset to movement offset) for the far target compared to the narrow target. Indeed, the movements towards the far targets were, on average, 320ms (SE: 24ms) longer than the movements towards the near target (main effect of target distance: F(1,29)=183.22, p<0.0001, partial eta-square: 0.86).

Similarly to movement curvature, the pattern of within-subject variability across movements in one condition was different for the near and far targets. While across-movement variability only slightly increased during the movement for the near target, there was a marked transient increase in across-movement variability for the far target. Movement variability peaked around the middle of movement before decreasing to levels similar to those for the near target close to the target (Fig.3). At the time of line bisection, variability was higher for the far target than for the near target (main effect of target distance: F(1,29)=19.4, p=0.00013, partial eta-square: 0.4). This effect was larger for the horizontal lines than the vertical ones (interaction between line orientation and target distance: F(1,29)=8.32, p=0.007; post-hoc Tukey test: H: p=0.0001; V: p= 0.3)

**Figure 3:**
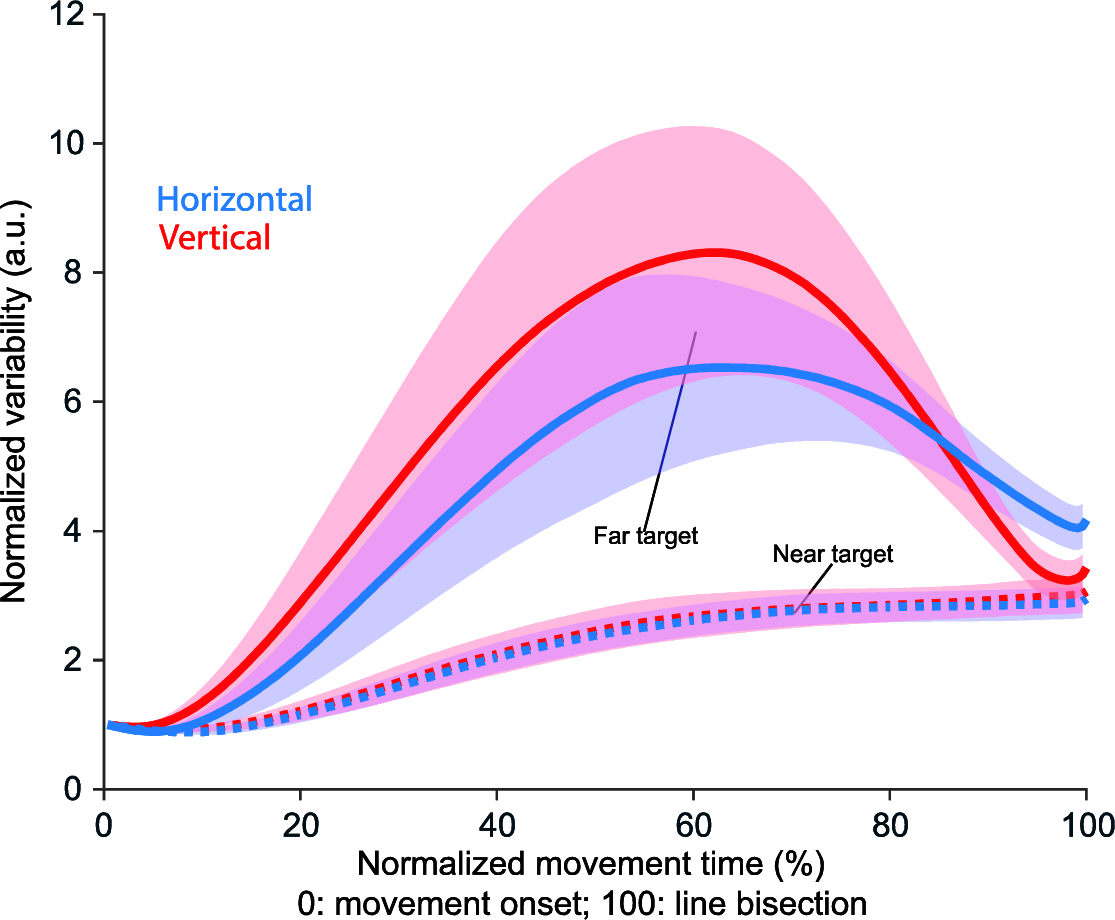
Influence of line orientation and distance on normalized within-subject variability across movements. Blue is associated with horizontal lines (H) and red with vertical lines (V). Solid lines are associated with the far targets and dotted ones with the near targets. Error bars represent 95% confidence interval.

There was a lot of variability in how the curvature of movements changed with target distance (compare Fig.1A and 1B). Therefore, we tested whether participants who exhibited large changes in movement curvature had a larger change in reaction time with target distance than people who performed straight movements towards the target independently of target distance (Fig.1B). To do so, we correlated the change in median reaction time from near to far targets with the change in movement curvature from near to far targets. This was performed independently for each line orientation (Fig.4). We found that the changes in curvature with movement distance was correlated with the change in reaction time with movement distance across subjects (H: r=-0.45, p=0.01; V: r=-0.54, p=0.002). It should be noted that reaction time and curvature for each distance and line orientation were not correlated, probably because of the large inter-individual differences in average reaction time. Similarly, the change in reaction time was uncorrelated with the change in absolute error at the end of the movement (X: r = −0.2263, p = 0.23; Y: r = 0.0333, p = 0.86) or the change in endpoint movement variability (X: r = −0.08, p = 0.66; Y: r = 0.2, p = 0.28).

**Figure 4:**
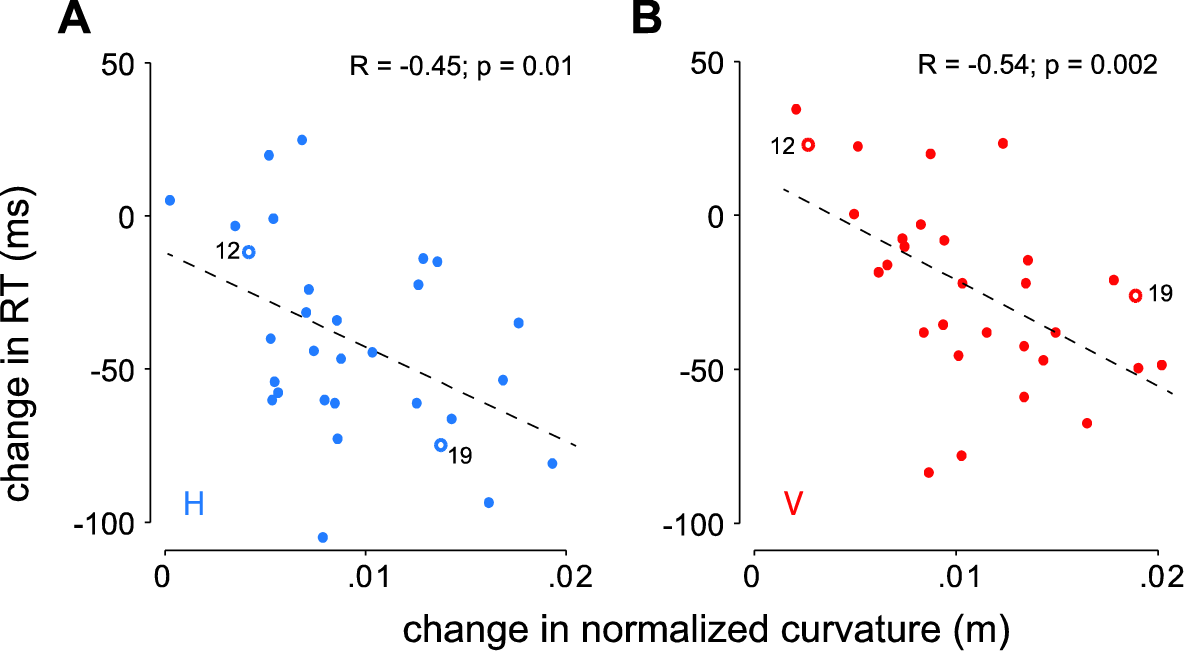
Relationship between change in mean normalized movement curvature for far to near targets and change in median reaction time for far to near targets across subjects. Left panel is for movements towards horizontal lines and right panel for vertical lines. In each panel, the two points that are highlighted (open circles) and numbered correspond to the values for the two subjects illustrated on Fig.1.

The change in target distance influenced movement curvature but also movement speed (main effect of target distance on movement peak velocity: F(1,29)=137.9, p<0.0001). In addition, peak velocity of the movement was correlated with reaction time. We observed that an increase in movement speed was coupled with an increase in reaction time (X: r=0.54, p=0.02; Y: r=0.35, p=0.057). In order to make sure that the effect of movement curvature on RT was independent of movement speed, we performed a partial correlation analysis and found that, even when the change in movement speed was taken into account, changes in movement curvature and changes in movement reaction time were still correlated (X: r=-0.38, p=0.038; Y: r=-0.48, p=0.007).

## Discussion

In this study, we showed that the reaction time of a reaching movement was affected by the chosen hand trajectory. Curved movements made toward a more distant target had, on average, a shorter reaction time than straighter movements made toward a nearer target. Furthermore, this effect depended on the curvature of the movement. A larger increase in movement curvature for the far compared to the near target was accompanied by a larger drop in movement reaction time. This decrease in reaction time was not correlated with movement accuracy or variability.

### Confounding factors

Movement complexity, movement extent or the presence of a sequence of movements or not differ between movements to near and far targets. Interestingly, all these possible confounding factors would result in an increase in reaction time for movements towards the far targets compared to the near targets (i.e. opposite to what was observed). For instance, several studies have shown that an increase in movement complexity is accompanied by longer reaction times (Henry and Rogers, 1960; Klapp et al., 1974; Christina and Rose, 1985). Similarly, larger movements are traditionally associated with longer reaction times (Munro et al., 2007; Falco et al., 2013). Finally, an increased number of movement phases, such as the ones we sometimes observed towards the far target, has been associated with an increase in reaction time (Klapp, 1995).

### Taking accuracy demands into account later

In contrast to Laszlo and Livesey (1977) who showed that the presence or absence of accuracy demands had a huge impact on reaction time, we observed a modulation of reaction time with target distance despite matched accuracy demands across conditions. In our study, while movements to the near targets had to take the accuracy demand into account before movement onset, the influence of accuracy demands for movements to the farther target could be delayed to later during the movement. As a result, reaction times were shorter for the farther target.

This hypothesis is consistent with the fact that movement variability increased during the first phase of the movements towards the far target (a sign that accuracy demands are not so important early in the movement) but was reduced during the second phase of the movement (Fig.3). This is compatible with the existence of two movement phases: a transport phase during which across-movement variability increased and a second phase during which movement variability is reduced. In contrast, movement variability remained much smaller during movements towards the near target (Fig.3), which point that the existence of these two phases is specific to movement to the farther target.

This hypothesis can also account for the correlation between movement curvature and reaction time (Fig.4). Following this hypothesis, highly curved movements (e.g. Fig.1A), which often consisted of two velocity peaks, are composed of two sub-movement parts: a first sub-movement that aimed at transporting the hand to an intermediate location while the second sub-movement was responsible for bringing the hand to bisect the target line as accurately as possible. In contrast, participants who moved their hand directly towards the far target (e.g. Fig.1B) did not exhibit such a transport phase and directly aimed at the center of the line. Therefore, for these movements, accuracy demands are taken into account from movement initiation on. This suggests that participants who postponed the processing of accuracy demands had a shorter reaction time for far targets compared to near targets than participants who did not. Similar effects of accuracy demands on reaction time have been found for catch-up saccades that are executed during smooth pursuit eye movements (Orban de Xivry and Lefèvre, 2007). That is, saccades that were updated online to integrate new information on the fly, had a shorter reaction time than the straighter saccades that were not modified during the movements (Schreiber et al., 2006). Together with our results, this suggests that the brain can either decide to start a movement early and to modify it later on the basis of new incoming information or that it can delay the onset of a movement and make it more straight (i.e. no change in goal during movement execution).

Importantly, our data highlight that taking accuracy demands into account in the planning of movement trajectory is not a black or white process as suggested by the study of Laszlo and Livesey (1977). Rather, we observed a continuum of change in reaction times. For straight hand trajectories to the far target, accuracy demands are taken into account before movement onset. For such straight movements towards the target, reaction time is not influenced by the loosening of accuracy demands (as in Orban de Xivry, 2013; Orban de Xivry and Lefevre, 2016) but just by their presence. Alternatively, delaying the effect of accuracy demands on hand trajectory (by refining movement plan during its execution), resulted in a gradual decrease in reaction time.

### Impact of these results on theories of motor planning

The reaction time is widely considered as an important variable to understand motor planning as it is considered that all the processes related to motor planning take place during that interval (Wong et al., 2015; Haith et al., 2016). Indeed, there is a widely accepted hypothesis that motor commands are assembled prior to movement execution during the motor planning stage. However, the present data reveal a different picture. Taking accuracy demands during motor planning seems to be a dynamic process with the amount of accuracy demands taken into account increasing over time. Independently of how loose they are, taking into account accuracy demands imposed a reaction time penalty. This penalty is reduced if the start of the movement can consist of a transport phase and therefore the accuracy demands can be incorporated into motor commands later in the movements, hence reducing the cost of accuracy on reaction time.

Such specification of motor commands prior to movement onset is also a hallmark of the state-space theory of motor planning where neurons in the motor cortex are thought to reach a given neural state before the movement is actually started (Shenoy et al., 2013; Ames et al., 2014; Kaufman et al., 2014). One can then wonder how the neural state looks like when motor preparation is not complete prior to movement onset but is further refined early during the movement. One prediction is that preparatory activity (in the null space following Kaufman et al., 2014) should be observed during movement execution. Our paradigm can provide information about how delaying the influence of accuracy demands on movement trajectory can influence the neural state of the motor cortex prior to and during movement.

## Conclusion

In this study, we demonstrate that the presence of accuracy demands for reaching movements greatly impacts reaction time. That is, reaction time can be up to 100ms longer in movements for which accuracy demands are immediately taken into account in comparison with movements for which the influence of accuracy demands on movement kinematics is delayed. In contrast to what current theories of motor planning suggest, these results show that movement kinematics can be refined during movement execution in order to reduce reaction time at movement initiation.

## Acknowledgement

We thank L. Van Leeuw for help with data collection and Tobias Heed for comments on an earlier draft of this manuscript. This work was supported by the Belgian Program on Interuniversity Attraction Poles and PRODEX initiated by the Belgian Federal Science Policy Office, Actions de Recherche Concertée (French community, Belgium) and the European Space Agency (ESA) of the European Union. VL is supported by the Research Fund of the French-speaking Community of Belgium (F.R.S.-FNRS). JJO, VL designed the experiment. JJO and L. Van Leeuw collected the data. JJO analysed data and drafted the paper. JJO, VL and PL finalized the paper.

